# Gastric epithelium from *BRCA1* and *BRCA2* carriers harbors increased double-stranded DNA damage and augmented growth

**DOI:** 10.1101/2025.10.01.679809

**Authors:** Kole H. Buckley, Michaela E. Dungan, Kevin Dinh, Gregory M. Kelly, Ryan Hausler, Kate E. Bennett, Daniel G. Clay, Julia E. Youngman, Ariana D. Majer, Keely A. Beyries, Blake A. Niccum, Sydney M. Shaffer, Tatiana A. Karakasheva, Kathryn E. Hamilton, Michael L. Kochman, Gregory G. Ginsberg, Nuzhat Ahmad, Kara N. Maxwell, Bryson W. Katona

## Abstract

An accumulating body of evidence suggests carriers of a pathogenic germline variant (PGV) in *BRCA1* or *BRCA2* have increased gastric cancer (GC) risk. *BRCA1* and *BRCA2* are tumor suppressor genes involved in promoting homologous recombination to repair double-stranded DNA breaks. The aim of this investigation was to identify differences within the gastric epithelium and in patient-derived gastric organoids (PDGOs) between *BRCA1* and *BRCA2* carriers and non-carriers to determine if evidence of early gastric carcinogenesis exists amongst these carriers. First, using gastric epithelial biopsies, *BRCA2* carriers were found to harbor higher expression of the proliferative marker Ki-67 within the antral gastric epithelium and strikingly, biopsies from both *BRCA1* and *BRCA2* carriers displayed a marked increase in double-stranded DNA damage. These results were further explored using PDGOs, where a growth advantage was observed for both *BRCA1* and *BRCA2* PDGOs compared to non-carrier PDGOs. Furthermore, both *BRCA1* and *BRCA2* PDGOs displayed a more pronounced enhancement of Ki-67 expression as well as increased double stranded DNA damage compared to non-carrier PDGOs. Importantly, none of the PDGOs showed signs of *BRCA1* or *BRCA2* loss of heterozygosity, potentially indicating a haploinsufficient phenotype. Taken together, these novel findings suggest that haploinsufficiency in *BRCA1* and *BRCA2* carriers may lead to DNA damage in the gastric epithelium, which may serve as an early event contributing to GC development.

## Introduction

Breast cancer susceptibility gene one (*BRCA1*) and two (*BRCA2*) are tumor suppressor genes that play a crucial role in the repair of double-stranded DNA breaks [1–4]. Specifically, BRCA1 and BRCA2 are involved in the homologous recombination (HR) pathway of double-stranded DNA break repair [1–4]. The HR pathway is a near error free method of DNA repair that involves recruitment of a homologous DNA template to resynthesize damaged DNA at the site of a break [2–4]. The other main pathway of double-stranded DNA break repair is non-homologous end joining (NHEJ), which involves the direct ligation of the damaged ends of DNA [2–4]. Importantly, NHEJ is a more error prone method of double-stranded break repair that can result in small nucleotide deletions, which can accumulate over time and promote genetic instability and tumorigenesis [2–4].

The loss of function of BRCA1 or BRCA2 due to a pathogenic germline variant (PGV) in the *BRCA1* or *BRCA2* gene respectively can lead to homologous recombination deficiency (HRD), where a cell cannot rely on HR to efficiently repair double-stranded DNA breaks [2–4]. Instead under these circumstances, cellular repair of double-stranded DNA breaks is biased towards the more error-prone NHEJ method [2–4]. Because of this, *BRCA1* and *BRCA2* carriers have a well-established increased risk of multiple cancers, most notably, breast, ovarian, prostate and pancreatic cancers [5–7]. Importantly, there has been a steady accumulation of clinical data suggesting that *BRCA2*, and to a slightly lesser extent *BRCA1,* carriers also have an increased risk of gastric cancer (GC) [1, 8, 9]. One particular study that looked at the absolute risk of GC to age 80 in over 3,000 *BRCA1* and over 2,000 *BRCA2* families from across five continents found *BRCA1* male and female carriers to have a 1.6% and 0.7% risk of GC, respectively, and both male and female *BRCA2* carriers having a 3.5% risk of GC [9]. Notably, the risk of GC in male *BRCA1* carriers and both male and female *BRCA2* carriers is 2-4-fold higher than the general US population lifetime risk of GC (0.8%) [10]. Furthermore, while the baseline incidence of GC is higher in east Asian populations compared to Western populations, one study examining a large cohort of Japanese *BRCA1* and *BRCA2* carriers found a 19-21% cumulative risk of GC to age 85, whereas non-carriers had a 4% risk [8]. It is unclear whether *BRCA1* and *BRCA2*-associated GC favors a particular subtype (i.e., diffuse versus intestinal), or if GC preferentially forms in a particular region of the stomach, as such granular details are typically underreported in the existing literature [1].

Despite a growing body of clinical data suggesting an increased risk of GC among *BRCA1* and *BRCA2* carriers, there is limited research investigating the underlying pathogenesis, and at this time no concrete mechanism of gastric carcinogenesis [1]. Therefore, we aimed to begin to understand the pathogenesis of GC among *BRCA1* and *BRCA2* carriers with the initial primary goal of identifying differences in markers of growth, proliferation, and DNA damage within the gastric epithelium of *BRCA1* and *BRCA2* carriers, as these factors may contribute to increased GC risk. To accomplish this, we utilized fresh endoscopically obtained gastric biopsies from *BRCA1* and *BRCA2* carriers without a history of GC as well as patient-derived gastric organoids (PDGOs) from both the gastric body and gastric antrum regions of the stomach and compared these to biopsies and PDGOs from individuals who do not harbor a *BRCA1* or *BRCA2* PGV (control). Thus, this study was aimed at identifying differences in the gastric epithelium in *BRCA1* and *BRCA2* carriers that may serve as early events or contributors to gastric carcinogenesis.

## Materials and Methods

### Acquisition of gastric biopsies

Gastric biopsies of the body and antrum were collected from individuals who were undergoing endoscopic ultrasound (EUS) and/or esophagogastroduodenoscopy (EGD) as part of their routine care from 05/27/2020 to 10/11/2023 at Penn Medicine. All individuals provided informed consent through an IRB-approved gastric biopsy collection protocol (IRB #842961). Eight non-targeted biopsies from each participant were collected from the gastric body and eight non-targeted biopsies were collected from the gastric antrum. Two biopsies from the body and antrum were utilized for paraffin embedding and sectioning, while remaining biopsies were utilized to generate PDGOs as detailed below. Gastric biopsies were collected from five *BRCA1* PGV carriers, five *BRCA2* PGV carriers, and five controls (with germline genetic testing confirming these individuals did not carry a *BRCA1* nor *BRCA2* PGV) for a total of 15 participants.

### Biopsy processing

From each participant, two biopsies from the gastric body and antrum were immediately immersed in 4% formalin and allowed to fix overnight at 4°C. Biopsies were then separately placed in cassettes and submitted to the University of Pennsylvania Molecular Pathology and Imaging Core for subsequent paraffin embedding and sectioning. Biopsies were sectioned in 10µm slices.

### Generating PDGOs

PDGOs were generated as previously described [11]. Briefly, gastric biopsies from either the gastric body or gastric antrum were washed and cut into small (1-2 mm) pieces. Biopsy pieces were then digested down to single cells using a combination of collagenase type 3 (Worthington; LS004182) and dispase II (Sigma-Aldrich; D4693-1G) for 30 minutes, followed by a 10-minute incubation with 0.25% EDTA-trypsin (Gibco; 25200-056). Single cells were extracted and mixed with Matrigel (Corning; 354234) to then allow plating of 50 µL Matrigel droplets into wells of a 24-well tissue culture plate. Tissue culture plates were then inverted and placed in a tissue culture incubator for 35 minutes to allow the Matrigel to polymerize into a 3-dimensional “dome” shape. Seeding of single cells were standardized to 10^5^ cells per well. Organoid media was changed every 2-3 days. The organoid media consisted of Advanced DMEM/F12 (Gibco; 12634-010), 50% v/v L-WRN conditioned media, 10mM HEPES (Invitrogen; 15630080), 1X Glutamax (Gibco; 35050-061), 100 U/mL Pen Strep (Gibco, 15140-122), N2 supplement (Invitrogen; 17502048), 1X B27 (Invitrogen; 17504044), 50ng/mL hEGF (Peptrotech; AF-100-15), 10nM Gastrin-l (Sigma Aldrich; G9145), 2µm Y-27632 (Sigma Aldrich; Y0503), 100ng/mL hFGF-10 (Peprotech; 100-26), and 2µm A83-01 (R&D Systems; 2939) [11].

### Embedding PDGOs

To embed PDGOs, the media was aspirated from six wells of the tissue culture plate and replaced with cold Cell Recovery Solution (Corning; 354253). The plate was then placed on a rocker for 35 minutes at 4°C. Next, the six Matrigel “domes” containing the organoids were broken up by gently aspirating and dispensing the Cell Recovery Solution on to the “domes” of each well. Organoids were then aspirated and transferred to a 15mL conical tube and centrifuged at 1400 xg for 3 minutes. After removing the supernatant, the organoids were then transferred to a 1.5mL tube containing 500µL of 1X PBS (Gibco; 10010023) and centrifuged at 2500 xg for 1 minute. The supernatant was then removed and 50µL of bacto-agar (BD; DF0140-15-4) was added to the organoids and gently mixed by pipetting repeatedly. The organoid/bacto-agar mix was then quickly dispensed onto a plastic petri dish in a “dome” shape and the bacto-agar allowed to solidify for 35 minutes at room temperature. Afterwards, 4% formalin was added to the dish and allowed to fix for 3 hours or overnight. Lastly, the organoid/bacto-agar “domes” were gently dislodged using forceps and placed in tissue cassettes for paraffin embedding and sectioning.

### Imaging PDGOs

PDGOs were imaged at 10-, 15-, and 20-days post initiation using a Keyence BZ-X800 Cell Imaging Microscope. The entire Matrigel “dome” of each well was imaged, taking care to capture the full depth via z-stack. Z-stack images were merged and displayed as a max projection.

### Image analysis

CellSens (Olympus) image analysis software was used to measure PDGO size and morphology on max projection images of the entire Matrigel “dome” at 10-, 15, and 20-days post initiation from biopsy tissue. Organoid size was measured as area (µm^2^) and morphology was measured using a sphericity function within the software [12]. Organoid number was manually counted on max projection images. Two to four wells per participant were analyzed and averaged for statistical comparisons in organoid size, number, and morphology. CellSens was also used to count the percent Ki-67+ nuclei, as well as the number of γ-H2AX and 53BP1 foci per nucleus in PDGOs and gastric biopsies. Quantification of the percent Ki-67+ nuclei consisted of an average of 1,727 nuclei (range 883 – 2,531) analyzed across two biopsies from each participant and gastric region, and an average of 554 nuclei (range 317 – 864) were analyzed across a minimum of 15 PDGOs for each participant. For quantification of γ-H2AX and 53BP1 foci per nucleus, an average of 841 nuclei (range 515 – 1,200) were analyzed across two biopsies from each participant and gastric region, and an average of 620 nuclei (range 445 – 1,053) were analyzed across a minimum of 15 PDGOs for each participant.

### Immunofluorescence

Immunostaining for Ki-67, γ-H2AX, and 53BP1 were performed using the same general protocol on formalin-fixed and paraffin embedded PDGOs and gastric biopsies. All PDGOs used for immunostaining were embedded between passage four and five. Briefly, tissue samples were deparaffinized via two 3-minute washes with xylenes and subsequently dehydrated and rehydrated with two 3-minute washes of 100% ethanol, 85% ethanol, 70% ethanol, and then 50% ethanol. Antigen retrieval was performed by immersing slides into a beaker containing citrate buffer (0.05% Tween-20, 10mM sodium citrate) and microwaving on high power for five minutes and left to stand at room temperature for 10 minutes. Slides were again microwaved for five minutes and left to cool at room temperature for 30 minutes. The tissue was then blocked for 30 minutes using a blocking buffer containing 10% goat serum, 0.1% triton-X, and 0.1% saponin in 1X PBS. Primary antibodies were diluted using the blocking buffer and applied for one hour (Ki-67 (Invitrogen; MA5-14520) 1:200, y-H2AX (Millipore; 05-636-1) 1:200, 53BP1 (Novus; NB100-904) 1:200). Samples were then washed twice for three minutes using PBST (0.05% tween-20, 1X PBS). Secondary antibodies were then applied for one hour (Alexa Fluor 488 (Invitrogen; A11008) 1:400, CY5 (Invitrogen; A10524) 1:200). The tissue was again washed with PBST twice for three minutes. Lastly, Fluromount G containing DAPI (SouthernBiotech; 0100-20) was applied to each slide prior to applying a cover slip. Slides were allowed to dry at room temperature overnight and stored long term at 4°C. Ki-67 immunostaining was imaged at 40x using a Leica Aperio Slide Scanner microscope. γ-H2AX and 53BP1 immunostaining were imaged at 60x using a Nikon Eclipse Ti-U widefield microscope.

### Comet assay

The neutral comet assay was performed to quantify double-stranded DNA breaks [13]. A comet assay kit (R&D Systems; 4250-050-K) was used according to the manufacturer’s instructions. Briefly, PDGOs between passage five and six were dissociated to 10^5^ single cells and mixed with LMAgarose (R&D Systems; 4250-050-02). Fifty µm of the cell-agarose mixture was then dispensed onto CometSlides (R&D Systems; 4250-050-03) and incubated in a lysis solution (R&D Systems; 4250-050-01) for one hour at 4°C. Slides were then immersed in 1X neutral electrophoresis buffer (100mM Tris Base, 300mM sodium acetate trihydrate) for 30 minutes at 4°C. Slides were then placed in an electrophoresis chamber with 4°C 1X neutral electrophoresis buffer and ran for 45 minutes at 21 V. Slides were removed and immediately immersed in a DNA precipitation solution (7.5 M ammonium acetate in 95% ethanol) for 30 minutes. Afterwards, slides were incubated for another 30 minutes in 70% ethanol. Slides were then allowed to dry for 20 minutes in a 37°C oven or until the agarose gel flattened on the slide. Lastly, slides were stained with SYBR Gold (Invitrogen; S11494) for 30 minutes, rinsed briefly with ddH2O, and then allowed to fully dry in a 37°C oven protected from light. The next day slides were imaged using an EVOS M700 fluorescent microscope (Invitrogen). A minimum of 50 comets per sample were analyzed via percent DNA in tail which was calculated as follows: 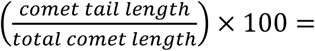 *percent DNA in tail* [13].

### DNA isolation, library preparation, and hybridization

PDGOs between passages six and seven were dissociated to single cells and used for DNA isolation from each PDGO line (minimum of 10^5^ cells per PDGO line). DNA was also obtained from a corresponding peripheral blood sample from the same participants from whom the PDGO lines were derived. One control PDGO sample was not included as a corresponding peripheral blood sample from the participant was unable to be obtained. Cellular DNA was isolated using a QIAmp DNA Mini Kit (Qiagen; 51304) according to the manufacturer’s protocol. The library was prepped for next generation sequencing (NGS) using the KAPA HyperPrep Kit (ROCHE; 7962363001) according the manufacturer’s recommended protocol and was hybridized using the xGen Hybridization and Wash Kit (IDT; 1080577) to a custom, targeted, and validated panel of 512 cancer genes [14, 15], as well as a spiked in xGen CNV Backbone Panel (IDT; 1080563). Samples were sequenced using an Illumina NovaSeq 6000 at the Center for Applied Genomics core facility at the Children’s Hospital of Philadelphia (CHOP).

### DNA sequencing analysis

FASTQ files from sequencing of 28 PDGOs (body-and antral-derived PDGOs from 14 participants), and 14 matched peripheral blood isolated DNA samples were aligned to human genome version 38 (hg38) using BWA-mem version 0.7.17 [16]. No samples were removed due to sequencing quality control. Germline variants were called from BAM files using Genome Analysis Toolkits (GATK) HaplotypeCaller version 3.7 [17]. Somatic copy number variations were called using both Sequenza version 3.0.0 [18] and CNVKit version 0.9.9 [19]. Using CNVKit copy number segments as input, HRDex version 0.0.0.9 [20] was used to determine homologous recombination deficiency (HRD) scores including segments of large state transitions (LSTs), genomic loss of heterozygosity (LOH) and non-telomeric allelic imbalance (nTAI). Copy number burden as a measure of overall chromosomal instability (CIN) was calculated as the percentage of bases measured that were in an altered copy number state. Somatic single nucleotide variants (SNVs) were called using Mutect2 version 4.1.2 [21]. Somatic SNVs were annotated using vep version 104.3 [22] and OncoKB [23]. Tumor mutational burden (TMB) was calculated as the number of non-oncogenic SNVs with a depth greater than 20 and an allelic balance greater than 0.05 divided by the number of bases captured and multiplied by 100000.

### Statistics

Data sets were analyzed using one-way and two-way analysis of variance (ANOVA) using GraphPad Prism. The Tukey’s multiple comparisons *post-hoc* test was used to locate differences when the observed F ratio was statistically significant (p < 0.05). Data are presented as mean ± SD.

### Data availability statement

The data generated in this study are available upon request from the corresponding author.

## Results

### Cohort characteristics

The cohort studied consisted of five control individuals (with germline genetic testing confirming no *BRCA1* or *BRCA2* PGV), five *BRCA1* carriers, and five *BRCA2* carriers (Table S1). All *BRCA1* and *BRCA2* PGV carriers were heterozygous for their respective PGVs. The mean age at biopsy acquisition was 63 (SD 9.8 years), 47% were female, and all were of self-reported White race. No participants had a personal history of GC, 47% had a personal history of a cancer other than GC, and one (6.7%) participant (*BRCA1* carrier) had a family history of GC. Lastly, none of the participants had a current or known prior *Helicobacter pylori* (*H. pylori*) infection.

### BRCA2 antral gastric biopsies exhibit enhanced Ki-67 expression

We first assessed proliferation markers for baseline differences in the gastric mucosa of *BRCA1* and *BRCA2* carriers compared to controls. Ki-67 immunolabeling was performed on gastric biopsies from both the body and antral regions of the stomach, (Figure 1A). No statistically significant differences in percent Ki-67+ nuclei were detected between groups amongst gastric biopsies from the body (Figure 1B). However, antral gastric biopsies from *BRCA2* carriers displayed increased Ki-67 expression compared to control (Figure 1C).

**Figure 1:**
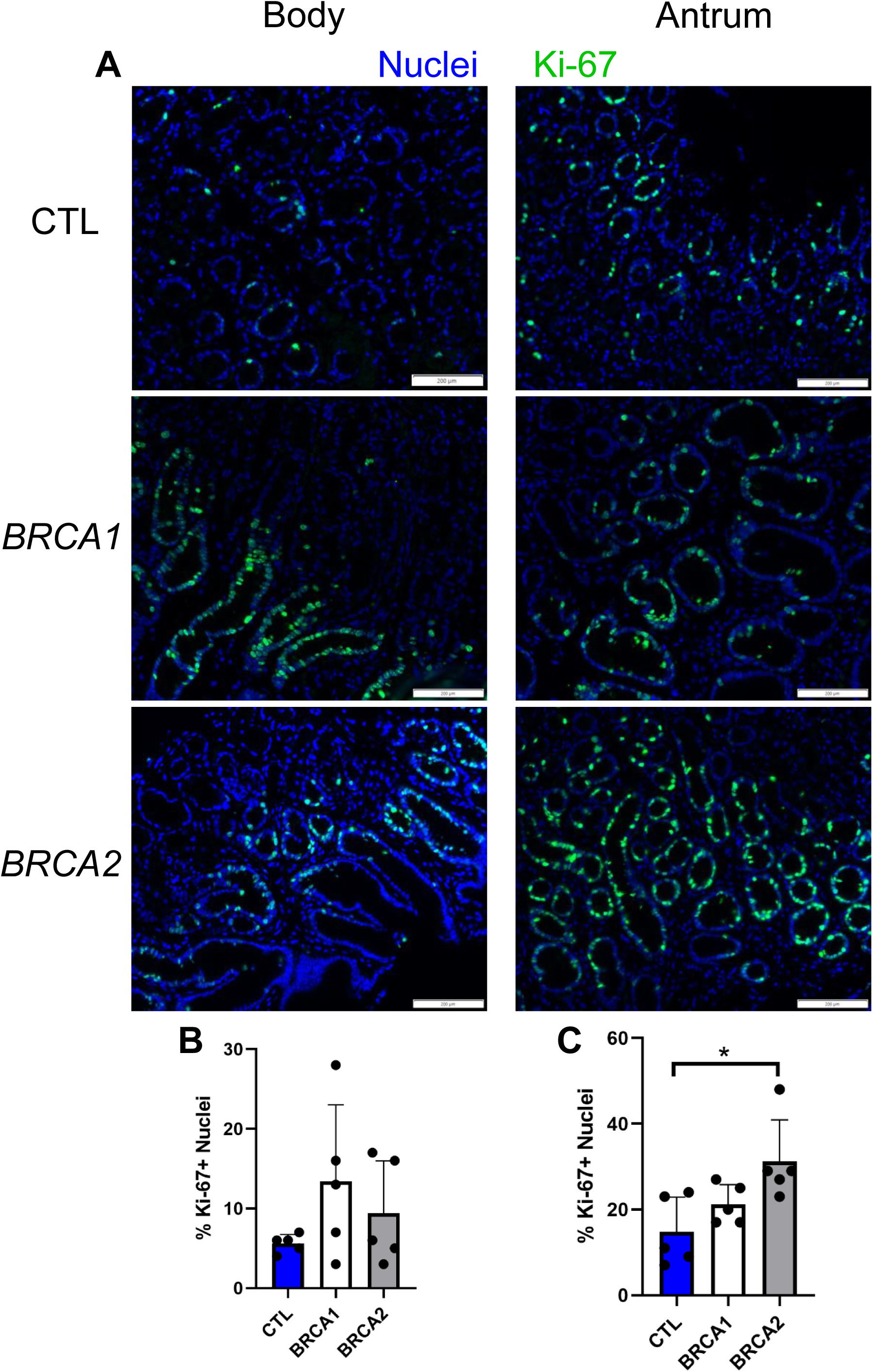
Antral biopsies from *BRCA2* carriers display enhanced proliferation. A) Representative images of Ki-67 immunolabeling for body and antral biopsies, scale bars = 200 µm. B) Percent Ki-67+ nuclei in body biopsies. C) Percent Ki-67+ nuclei in antral biopsies. * = p-value < 0.05 compared to indicated group. CTL = Control. *n* = five participants per group.

### BRCA1 and BRCA2 gastric biopsies display evidence of increased DNA damage

Given BRCA1 and BRCA2’s role in DNA double-stranded break repair, we next sought to uncover if there were differences in the prevalence of double-stranded DNA damage within the gastric mucosa of *BRCA1* and *BRCA2* carriers. We assessed double-stranded DNA damage via immunolabeling for two markers of DNA double-stranded breaks, γ-H2AX and 53BP1 [24] (Figure 2A-B).

**Figure 2:**
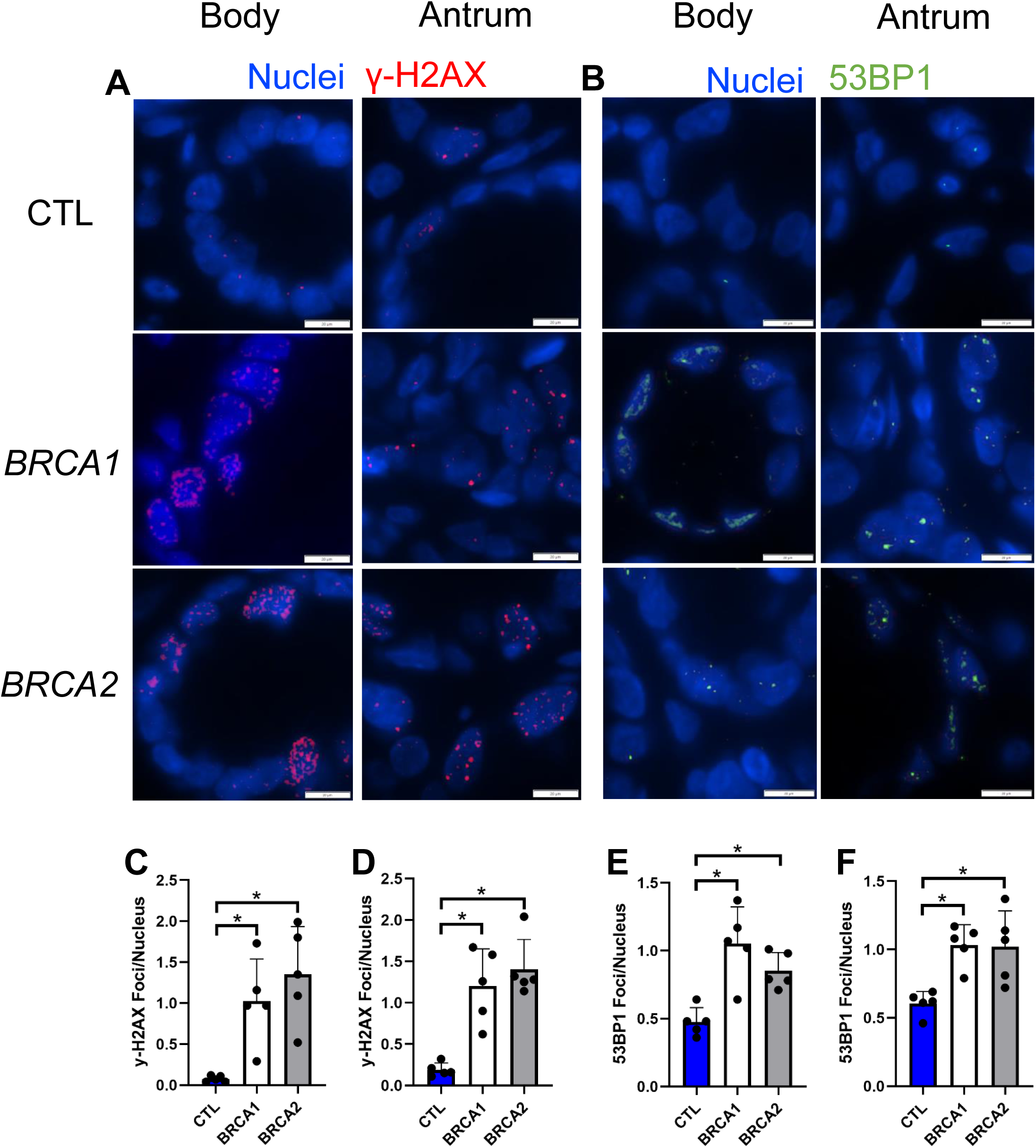
Body and antral biopsies from *BRCA1* and *BRCA2* carriers have increased double-stranded DNA damage. A) Representative images of γ-H2AX immunolabeling of body and antral biopsies, scale bars = 20 µm. B) Representative images of 53BP1 immunolabeling of body and antral biopsies, scale bars = 20 µm. C) Number of γ-H2AX foci per nucleus in body biopsies. D) Number of γ-H2AX foci per nucleus in antral biopsies. E) Number of 53BP1 foci per nucleus in body biopsies. F) Number of 53BP1 foci per nucleus in antral biopsies. * = p-value < 0.05 compared to indicated group. CTL = Control. *n* = five participants per group.

Both body and antral biopsies of *BRCA1* and *BRCA2* carriers showed on average an 11-fold greater prevalence of γ-H2AX foci per nucleus compared to controls (Figure 2C-D). Likewise, both *BRCA1* and *BRCA2* body and antral biopsies showed on average a nearly two-fold higher number of 53BP1 foci per nucleus compared to controls (Figure 2E-F). Taken together, these findings suggest there is increased prevalence of DNA damage throughout the gastric epithelium of both *BRCA1* and *BRCA2* PGV carriers.

### BRCA1 and BRCA2 PDGOs exhibit augmented growth

To further investigate the findings from *BRCA1* and *BRCA2* biopsy samples, we next sought to assess for differences in growth, morphology, proliferation, and DNA damage using PDGOs. Importantly, our previous work as well as the work of others has shown differences in growth between PDGOs generated from the gastric body and the gastric antrum [11, 25]. Therefore, body and antrum PDGOs were separately generated and analyzed.

We first aimed to assess the impact of *BRCA1* and *BRCA2* on organoid morphology. Day 20 representative images of body and antrum PDGO growth from a control participant, as well as a *BRCA1* and *BRCA2* carrier, are shown in Figure 3A. On average, there were no observed differences in body nor antral PDGO morphology as measured by sphericity (where a value of one equals a perfect sphere; Figure 3B-C) [12], suggesting that *BRCA1* and *BRCA2* do not play a role in PDGO morphology.

**Figure 3:**
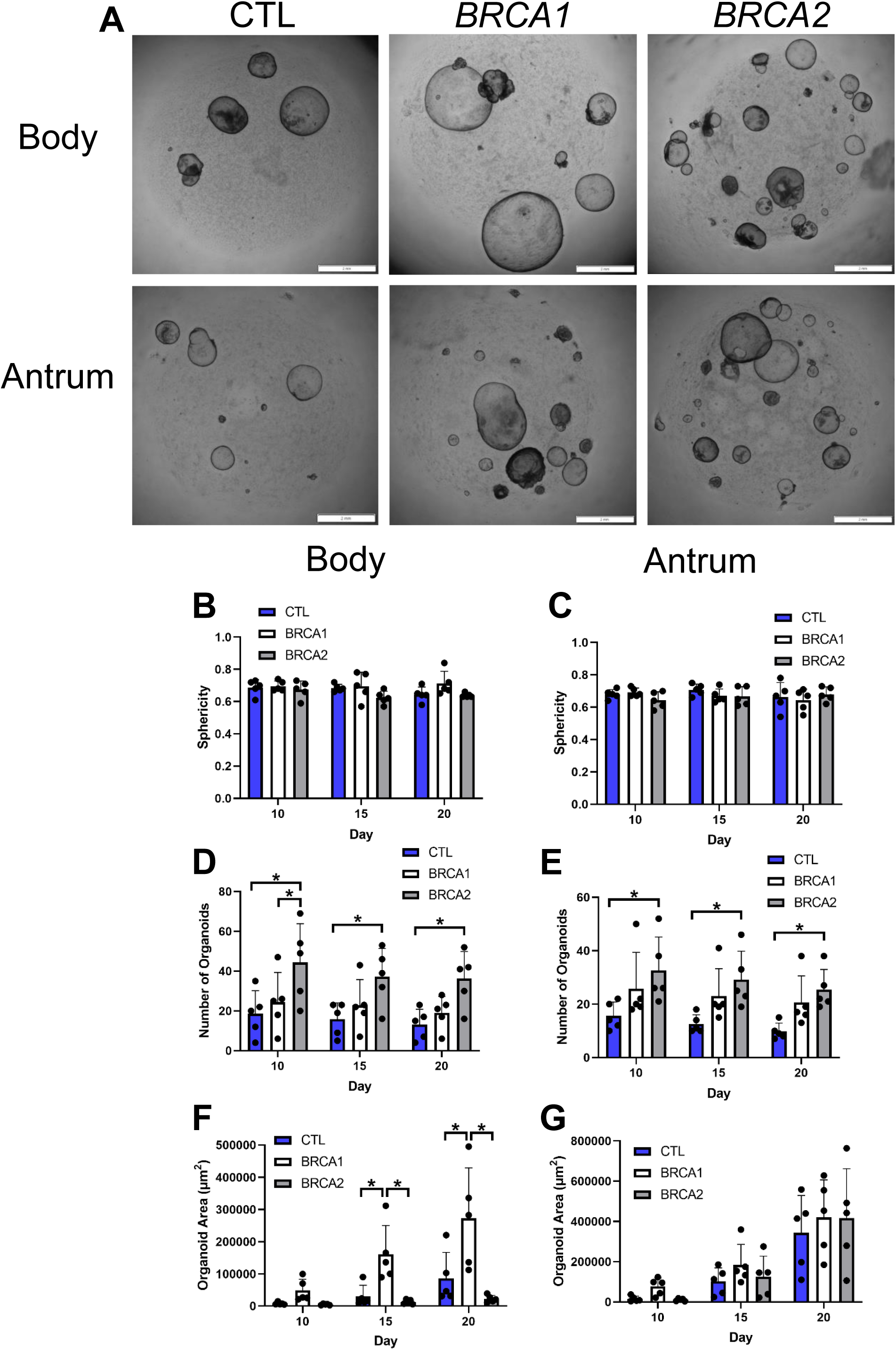
Growth differences amongst *BRCA1* and *BRCA2* PDGOs. A) Representative z-projection images of body and antral-derived PDGOs 20 days post initiation, scale bars = 2 mm. B) Morphology of body-derived PDGOs as measured via sphericity (where a value of one equals a perfect sphere). C) Morphology of antral-derived PDGOs as measured via sphericity (where a value of one equals a perfect sphere). D) Average number of organoids formed per well in body-derived PDGOs. E) Average number of organoids formed per well in antral-derived PDGOs. F) Average size of body-derived PDGOs per well as measured via area (µm^2^). G) Average size of antral-derived PDGOs per well as measured via area (µm^2^). * = p-value < 0.05 compared to indicated group, at indicated timepoint. CTL = Control. *n* = five participants per group.

Next, we assessed whether there were differences in the number of organoids formed at 10-, 15-, and 20-days after uniform initiation with 10^5^ gastric biopsy-isolated cells. Indeed, the number of organoids formed from *BRCA2* carriers was over 2-fold greater compared to controls at all time points for both body (Figure 3D) and antral (Figure 3E) PDGOs. While the number of organoids formed from *BRCA1* carriers trended toward being higher than controls at all time points for both body and antral PDGOs, no statistically significant differences were observed. We then sought to determine if there were differences in the average size of individual organoids as measured via area (µm^2^). For body PDGOs (Figure 3F), individual *BRCA1* organoids were on average over 2-fold larger than both controls and *BRCA2* carriers at 15-, and 20-days post initiation. No differences in size were observed in PDGOs generated from the antrum (Figure 3G). Taken together, we show an interesting dichotomy whereby body and antral *BRCA2* PDGOs have a higher formation rate while body *BRCA1* PDGOs grow larger in size.

### BRCA1 and BRCA2 PDGOs display enhanced Ki-67 expression

We next quantified the percentage of Ki-67+ nuclei in body and antral PDGOs (Figure 4A). Both *BRCA1* and *BRCA2* body PDGOs showed over a 3-fold higher percentage of Ki-67+ nuclei compared to controls (Figure 4B). For antral PDGOs, *BRCA2* organoids displayed nearly a 4-fold greater percentage of Ki-67+ nuclei compared to controls (Figure 4C).

**Figure 4:**
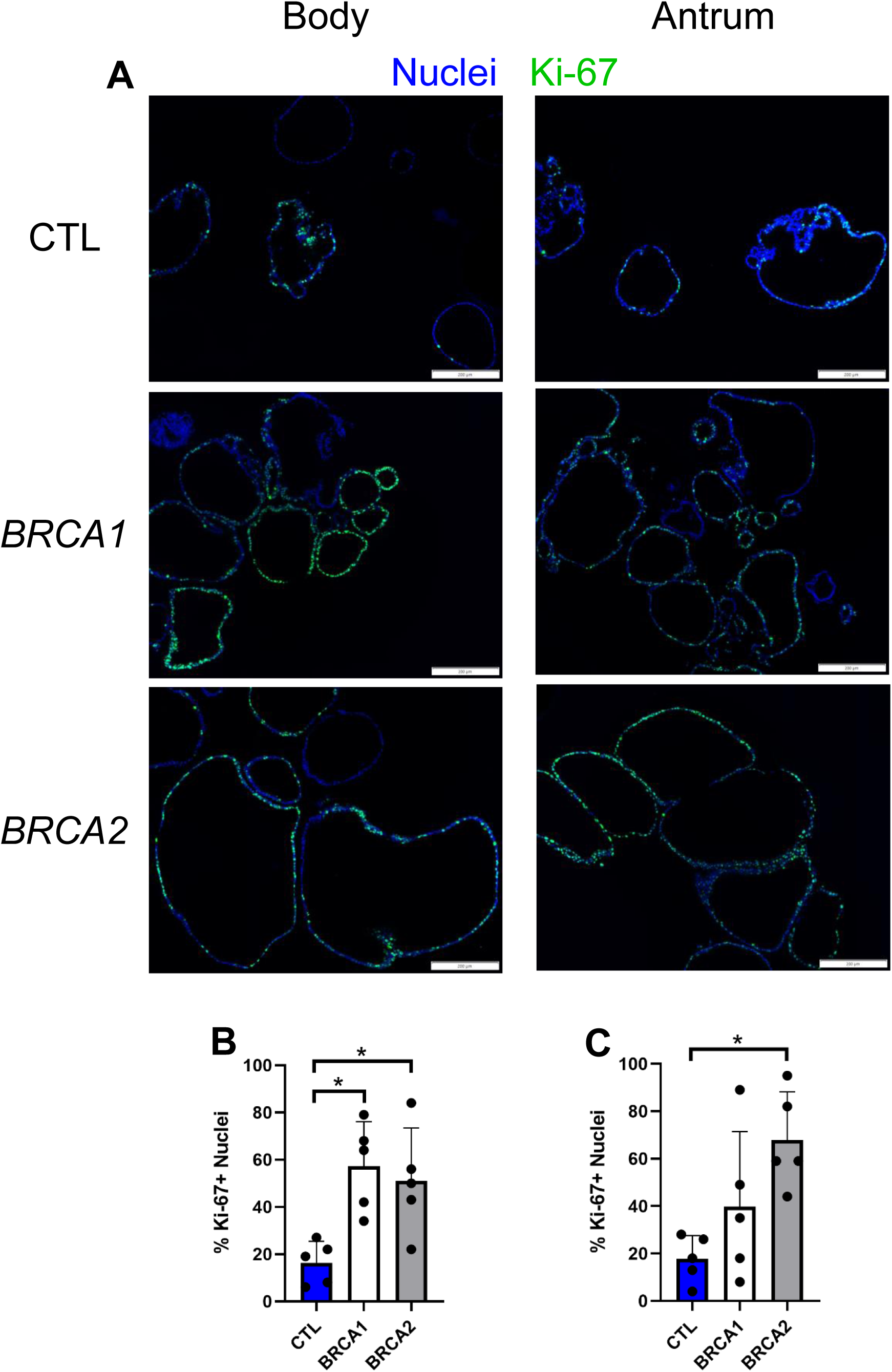
*BRCA1* and *BRCA2* PDGOs exhibit augmented proliferation. A) Representative images of Ki-67 immunolabeling of body and antral-derived PDGOs, scale bars = 200 µm. B) Number of Ki-67+ nuclei in body-derived PDGOs. C) Number of Ki-67+ nuclei in antral-derived PDGOs. * = p-value < 0.05 compared to indicated group. CTL = Control. *n* = five participants per group.

### BRCA1 and BRCA2 PDGOs show increased signs of double-stranded DNA damage

Given that gastric biopsies from *BRCA1* and *BRCA2* carriers showed increased DNA damage (Figure 2), we next assessed the prevalence of double-stranded DNA damage in *BRCA1* and *BRCA2* body and antral PDGOs (Figure 5A-B). The number of γ-H2AX foci per nucleus for both body and antral *BRCA1* and *BRCA2* PDGOs were on average 6.5-fold elevated over controls (Figure 5C-D). While no statistically significant differences were observed in the number of 53BP1 foci per nucleus for body PDGOs (Figure 5E), both *BRCA1* and *BRCA2* antral PDGOs had on average 1.8-fold more foci of 53BP1 per nucleus compared to controls (Figure 5F). The increased prevalence of double-stranded DNA damage in *BRCA1* and *BRCA2* PDGOs observed via y-H2AX and 53BP1 immunolabeling was indeed consistent with our findings from the corresponding gastric biopsies.

**Figure 5:**
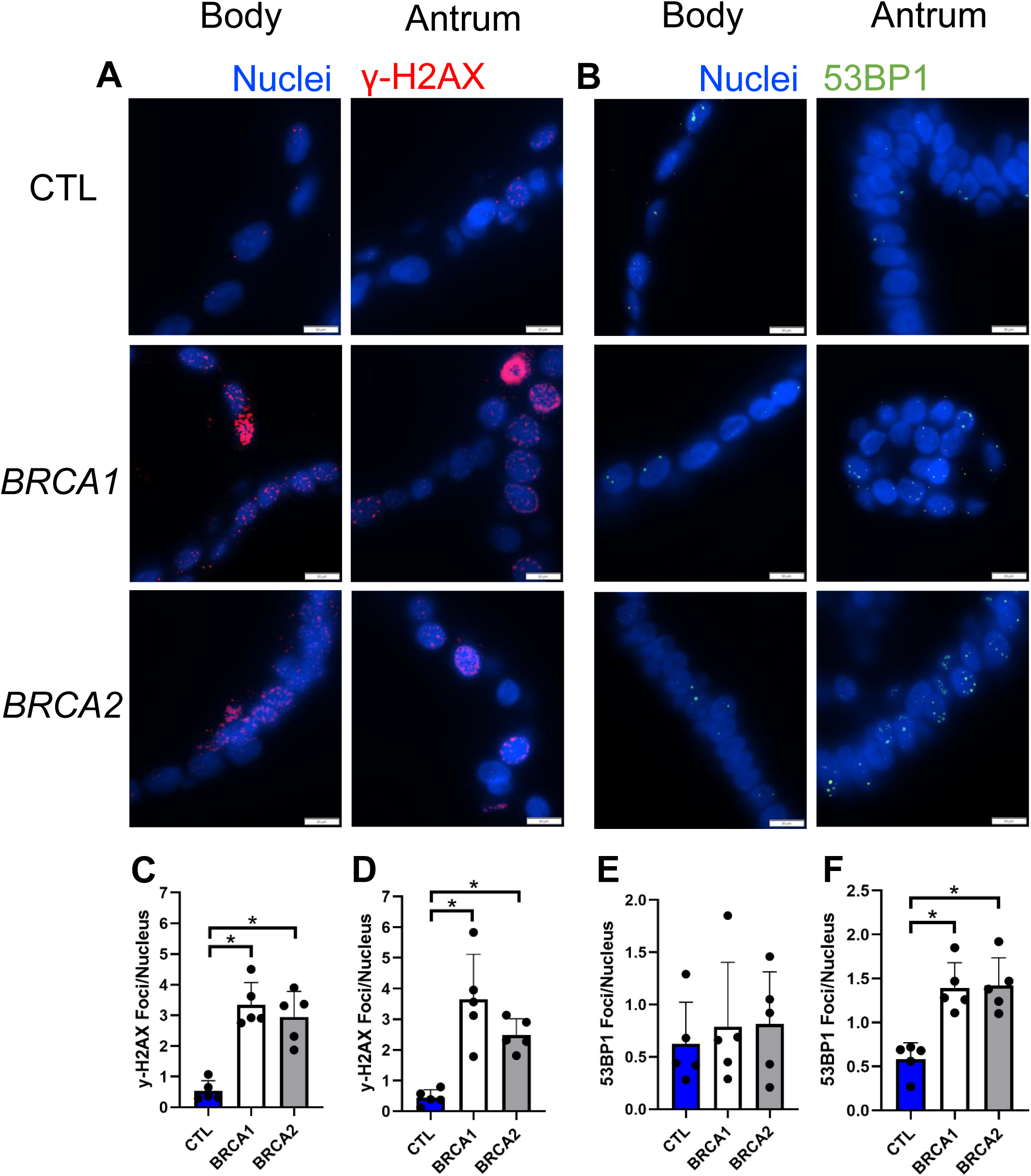
*BRCA1* and *BRCA2* PDGOs display increased double-stranded DNA damage. A) Representative images of γ-H2AX immunolabeling in body and antral-derived PDGOs, scale bars = 20 µm. B) Representative images of 53BP1 immunolabeling in body and antral-derived PDGOs, scale bars = 20 µm. C) Number of γ-H2AX foci per nucleus in body-derived PDGOs. D) Number of y-H2AX foci per nucleus in antral-derived PDGOs. E) Number of 53BP1 foci per nucleus in body-derived PDGOs. F) Number of 53BP1 foci per nucleus in antral-derived PDGOs. * = p-value < 0.05 compared to indicated group. CTL = Control. *n* = five participants per group.

To complement this immunolabeling technique with a secondary method of quantifying double-stranded DNA damage in PDGOs we employed the neutral comet assay. Specifically, we quantified the percent DNA in the comet tail as a readout of DNA double-stranded breaks, where a higher percentage of DNA in the comet tail represents a higher prevalence of double-stranded DNA damage [13] (Figure 6A). For body PDGOs, both *BRCA1* and *BRCA2* showed a 1.4-fold increase in the percentage of DNA in the comet tail compared to controls (Figure 6B). Likewise, antral PDGOs of both *BRCA1* and *BRCA2* carriers displayed on average a 1.5-fold increase in the percentage of DNA in the comet tail compared to controls (Figure 6C). Taken together, our double-stranded DNA damage findings in PDGOs and biopsies from *BRCA1* and *BRCA2* carriers suggest that *BRCA1* and *BRCA2* carriers harbor increased DNA damage within the gastric epithelium.

**Figure 6:**
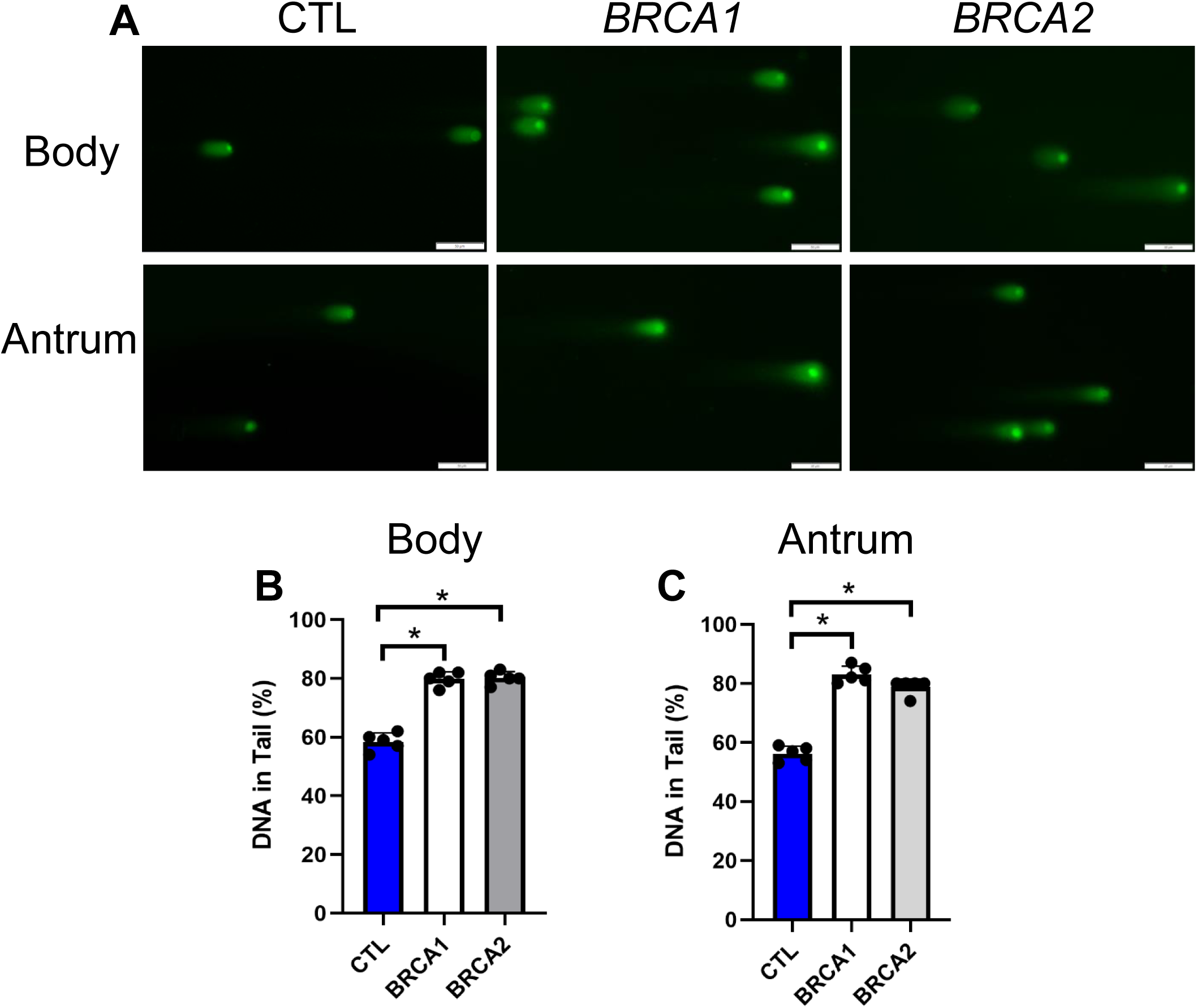
Comet assay demonstrates increased double-stranded DNA damage in *BRCA1* and *BRCA2* PDGOs. A) Representative images from the comet assay performed in both body and antral-derived PDGOs, scale bars = 50 µm. B) Percent DNA in comet tail quantification for body-derived PDGOs. C) Percent DNA in comet tail quantification for antral-derived PDGOs. * = p-value < 0.05 compared to indicated group. CTL = Control. *n* = five participants per group.

### BRCA1 and BRCA2 PDGOs show no loss of heterozygosity (LOH) or differences in genomic stability

Next, to determine if *BRCA1* or *BRCA2* LOH may have occurred leading to an increase in DNA damage, as well as to determine if increased DNA damage led to changes in genomic stability, we performed DNA sequencing on PDGOs and matched peripheral blood samples. Variant allele frequency (VAF) was determined and compared between DNA from a matched peripheral blood sample and the participant’s body and antrum PDGOs. VAF percentages ranged from 43 – 52% and allele-specific copy number data indicated a diploid heterozygous state of the *BRCA1* and *BRCA2* locus (Table S2), [26], indicating that none of the PDGOs used in our experiments underwent LOH (Figure 7A). Chromosomal instability (CIN) scores showed no significant differences between groups (Figure 7B). Mutational burden (MB), as measured by the number of genetic mutations per megabase (Mut/Mb), showed no significant differences between *BRCA1*, *BRCA2*, and control PDGOs (Figure 7C), nor did any samples reach a threshold for what would be considered a high MB [27]. Furthermore, homologous recombination deficiency (HRD) score, as measured by the unweighted sum of LOH, large-scale transition states (LST), and telomeric allelic imbalance (nTAI) scores, revealed no differences between *BRCA1*, *BRCA2*, and control PDGOs (Figure 7D-G), with only one *BRCA2* PDGO sample meeting the threshold for HR-deficiency [28, 29]. Taken together, the *BRCA1* and *BRCA2* PDGOs utilized in this investigation do not have LOH to explain the increased double stranded DNA damage or any major differences in genomic stability, compared to control PDGOs, indicating that the differences observed in this investigation are due to *BRCA1* and *BRCA2* haploinsufficiency.

**Figure 7:**
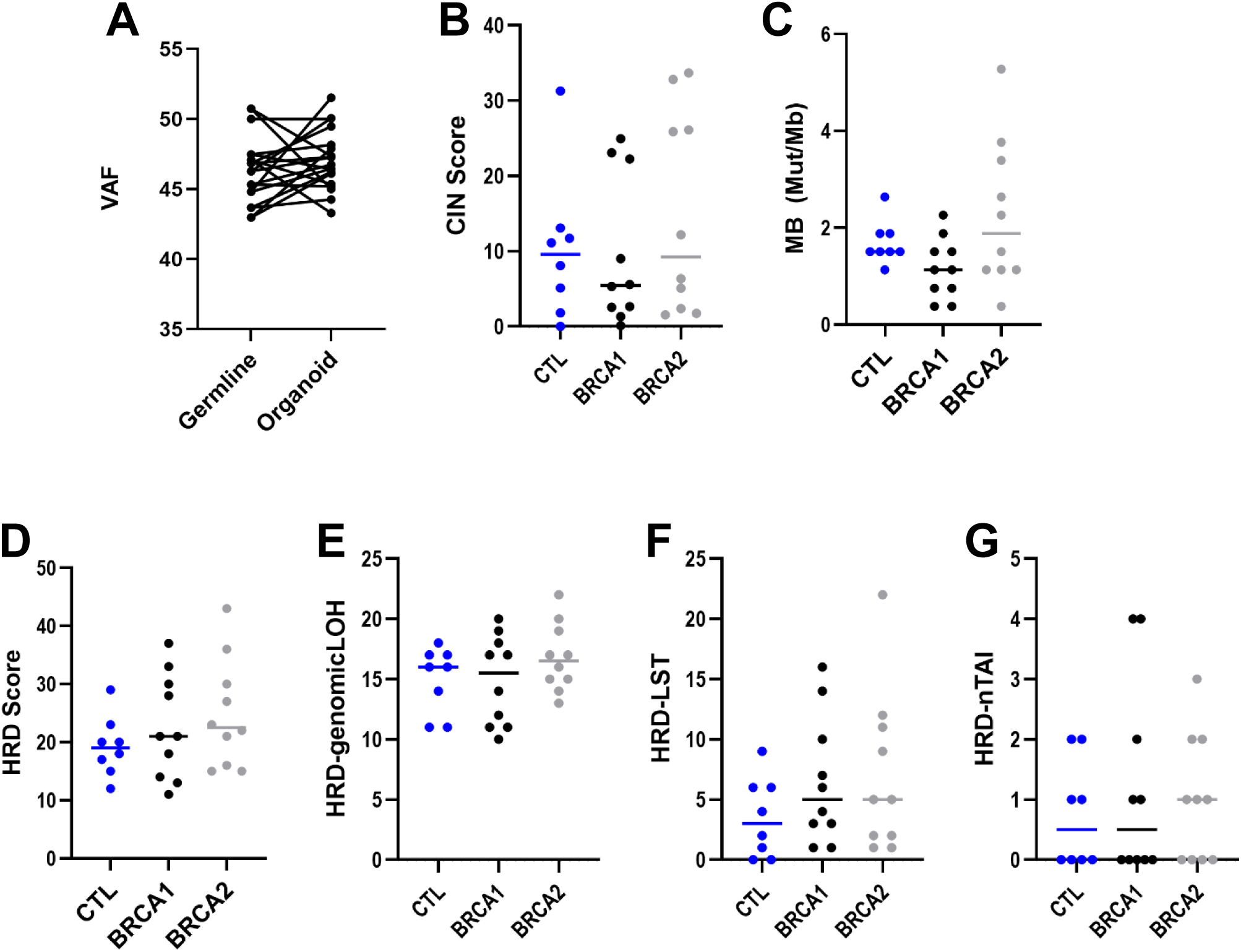
*BRCA1* and *BRCA2* PDGOs are genomically stable. A) Variant allele frequency (VAF) between all PDGOs and DNA from the corresponding patient’s blood sample. B) Chromosomal instability score (CIN) for body and antral PDGOS. C) Mutational burden (MB) as measured via the number of genetic mutations per megabase (Mut/Mb) for body and antral PDGOs. D) Homologous recombination deficiency (HRD) score based on the unweighted sum of E) loss of heterozygosity (HRD-genomicLOH), F) large-scale transition states (HRD-LST), and G) telomeric allelic imbalance (HRD-nTAI) scores of body and antral PDGOs. CTL = Control.

## Discussion

*BRCA1* and *BRCA2* carriers have a well-established risk of multiple cancers including breast, ovarian, prostate and pancreatic cancers [7]. In addition to these established cancer risks, a growing body of clinical data suggests *BRCA1* and *BRCA2* carriers also have an increased risk of GC [1]. However, at this time there is limited data exploring the pathogenesis of GC among these carriers, and the mechanism(s) of *BRCA1-* and *BRCA2*-associated gastric carcinogenesis remain uncertain. Therefore, the aim of this study was to utilize *BRCA1* and *BRCA2* carrier gastric biopsies and PDGOs to determine if the gastric epithelium of *BRCA1* and *BRCA2* carriers harbors differences from non-carriers, including changes in growth and/or DNA damage, that may begin to explain the increased risk of GC among these carriers, with the ultimate goal of informing future mechanistic studies into gastric carcinogenesis in *BRCA1* and *BRCA2* carriers.

To determine if the gastric epithelium of *BRCA1* and *BRCA2* carriers differed from non-carriers, we first assessed for proliferative markers and double-stranded DNA breaks in gastric epithelial biopsies from carriers as well as controls. For these initial studies we chose to examine biopsies from both the body and antrum separately as it remains uncertain if *BRCA1-* and *BRCA2*-associated GC preferentially forms in the proximal or distal stomach. Biopsies from the gastric antrum of *BRCA2* carriers displayed increased Ki-67 expression and both *BRCA1* and *BRCA2* carriers showed a striking elevation in markers of double-stranded DNA breaks (γ-H2AX and 53BP1) in both the body and antrum. Thus, our data suggests that there may be increased double-stranded DNA damage throughout the gastric epithelium of *BRCA1* and *BRCA2* carriers rather than having DNA damage localized to a particular region of the stomach.

These initial studies were performed on sections from endoscopically acquired gastric mucosal biopsies. While gastric mucosal biopsies are useful to study epithelial biology, there are many cell types present in gastric mucosal biopsies other than epithelial cells. Therefore, to study gastric epithelial proliferation and double-stranded DNA damage in a more pure epithelial model we then generated PDGOs from *BRCA1* and *BRCA2* carrier biopsies of both the gastric body and antrum. These organoid models showed more pronounced differences in Ki-67 expression and double-stranded DNA damage. We also found that both *BRCA1* and *BRCA2* carrier PDGOs showed signs of increased growth where *BRCA1* PDGOs grew larger, while *BRCA2* PDGOs formed in greater numbers compared to control PDGOs. With the increased growth in size of *BRCA1* PDGOs we evaluated the number of cells expressing Ki-67 for both gastric body and antral *BRCA1* PDGOs and observed a marked increase compared to controls. Furthermore, we also observed enhanced Ki-67 expression in *BRCA2* PDGOs generated from both the gastric body and antrum. Augmented proliferation is expected in GC-derived PDGOs [30, 31], and therefore it is possible that the augmented Ki-67 expression observed in *BRCA1* and *BRCA2* PDGOs may confer increased GC risk. Whether the enhanced Ki-67 expression observed is noted equally amongst all cells or is confined to a particular subpopulation of gastric epithelial cells remains uncertain and is an area of active investigation.

Use of PDGO models also allowed for confirmation that double-stranded DNA damage was observed at higher levels in the PDGOs from *BRCA1* and *BRCA2* carriers. Indeed, body and antral PDGOs from both *BRCA1* and *BRCA2* carriers showed robust signs of increased double-stranded DNA damage compared to control PDGOs via two different assays, including the comet assay that directly assesses for double-stranded DNA breaks. Studies investigating the prevalence of double-stranded DNA damage in the context of the gastric epithelium and/or GC are limited. While none of the participants in our cohort had a personal history of GC, there are a few studies that suggest increased double-stranded DNA damage in malignant gastric lesions [32–34] as well as in cases of early-onset GC [35]. In particular, one recent study noted a steady increase in the prevalence of double-stranded DNA damage in patient biopsies starting from normal gastric epithelium to gastritis and progressive stages of intestinal metaplasia, with the highest incidence noted in GC biopsies [34]. This data may indicate that increasing levels of DNA damage correlates with GC risk, which is noteworthy as we observed varying levels of DNA damage both in PDGOs and biopsies of *BRCA1* and *BRCA2* carriers. Whether or not higher levels of baseline DNA damage in the gastric epithelium is predictive of a *BRCA1* or *BRCA2* carrier’s risk of GC will need to be explored in future investigations.

The increased incidence of double-stranded DNA damage observed in both biopsies and PDGOs from *BRCA1* and *BRCA2* carriers is particularly concerning, as this type of damage is the most severe form of DNA damage and thought to be a general risk factor for cancer development [2, 36, 37]. Repeated insult to cellular DNA increases the likelihood of a cell acquiring pro-oncogenic mutations [2, 36, 37]. Additionally, HRD further heightens this risk, as DNA repair becomes reliant on the more error-prone NHEJ pathway [2–4]. It is through HRD that *BRCA1* and *BRCA2* carriers are thought to be at an increased risk for various cancers [5, 6]. However, DNA sequencing showed that only one of the PDGOs utilized in our investigation displayed signs of HRD. Furthermore, no differences in CIN and MB were observed when compared to control PDGOs. These findings were largely expected as none of the carriers utilized in this investigation had active GC nor a personal history of GC and the biopsies obtained were from normal appearing epithelium. It is possible that the increased levels of baseline DNA damage observed in *BRCA1* and *BRCA2* carriers acts a primer for gastric carcinogenesis, which upon addition of a second insult, such as a *Helicobacter pylori* (*H. pylori*) infection, may then more aggressively drive gastric carcinogenesis. *H. pylori* infects the gastric epithelium and has been well-established to increase GC risk by as much as 6-fold in the general population [38]. Strikingly, a recent study showed that *BRCA1* and *BRCA2* carriers (among other HR-related PGV carriers) with a concurrent *H. pylori* infection had up to a nine-fold increase in GC risk compared to uninfected carriers [39]. *H. pylori* itself is known to induce double-stranded DNA damage in gastric epithelial cells [40–42], which when coupled with a *BRCA1* or *BRCA2* carrier’s elevated baseline gastric epithelial double-stranded DNA damage, may result in a synergistic increase in double-stranded DNA damage of the gastric epithelium and promote the development of GC. This is an area under active investigation in our laboratory. Beyond *H. pylori*, a second insult to the gastric epithelium could come from other common sources of DNA damage in the gastric epithelium such as alcohol consumption and tobacco smoking, which are well-established risk factors of gastric cancer [43, 44]. It is possible that excessive alcohol consumption and/or tobacco smoking could similarly confer a synergistic increase GC risk in *BRCA1* and *BRCA2* carriers.

Importantly the *BRCA1* and *BRCA2* PDGOs utilized in this investigation displayed no signs of LOH. Although controversial, a growing body of literature suggests that LOH in *BRCA1* and *BRCA2* is not always necessary for tumorigenesis and instead *BRCA1* or *BRCA2* haploinsufficiency can promote tumorigenesis [45, 46]. One study utilizing normal primary breast epithelial cells derived from heterozygous *BRCA1* carriers noted enhanced proliferation despite no LOH [47]. The authors speculate that this elevation in proliferation may be a result of upregulation in the epidermal growth factor pathway (EGFR) [47]. Furthermore, other studies have noted increased DNA damage and/or DNA damage susceptibility in normal primary breast epithelial cells derived from *BRCA1* and *BRCA2* heterozygous carriers without LOH, which may be due to stalled replication forks [48–51].

### Limitations

This study has several limitations. The cohort used in this study is primarily older individuals (mean = 63 years old) and lacks racial and ethnic diversity. Additionally, none of the carriers in our study had active gastric cancer. Future investigations aimed at identifying a mechanism of carcinogenesis in *BRCA1* and *BRCA2*-associated GC may require the recruitment of carriers with active GC. Furthermore, additional studies with a larger cohort may be required to confirm that the differences observed in the 15 participants of the present study are more widespread across *BRCA1* and *BRCA2* carriers.

## Conclusion

To our knowledge, this is one of the first molecular studies undertaken to begin to understand *BRCA1* and *BRCA2*-associated GC risk among carriers. Taken together, our novel findings show that the gastric epithelium of *BRCA1* and *BRCA2* carriers harbor enhanced double-stranded DNA damage and augmented Ki-67 expression in the heterozygous state which suggests that these may be early events that contribute to GC risk in these carriers. Importantly, none of the PDGOs used in these experiments displayed signs of LOH or widespread genomic instability, indicating that BRCA1/BRCA2 haploinsufficiency may be contributing to the observed findings. Ultimately, a better understanding of gastric carcinogenesis in *BRCA1* and *BRCA2* carriers will have important implications for gastric cancer risk management amongst this high-risk cohort.

## Author Contributions

Conceptualization, KHB and BWK; Investigation, KHB, MED, KD, GMK, RH, and BWK. Resources, ADM, JEY, KEB, DGC, GMK, RH, BAN, SMS, TAK, KEH, KNM, MLK, GGG, NA, and BWK. Data curation, KHB, GMK, RH, KNM, and BWK, Writing -original draft, KHB and BWK. Writing -review and editing, all authors contributed. All authors have read and agreed to the published version of the manuscript.

## Acknowledgments

The authors would like to acknowledge support from the following: the NIH/NIDDK Center for Molecular Studies in Digestive and Liver Diseases at the University of Pennsylvania (P30DK050306), including the Molecular Pathology and Imaging Core facility, University of Pennsylvania Genomic Medicine T32 HG009495 (KHB), Men & BRCA Program at the Basser Center for BRCA (KHB, SMS, KNM, BWK), and the CHOP Gastrointestinal Epithelium Modeling Program and Core (RRID: SCR 026402) (TAK, KEH).

---

**Table S1.**
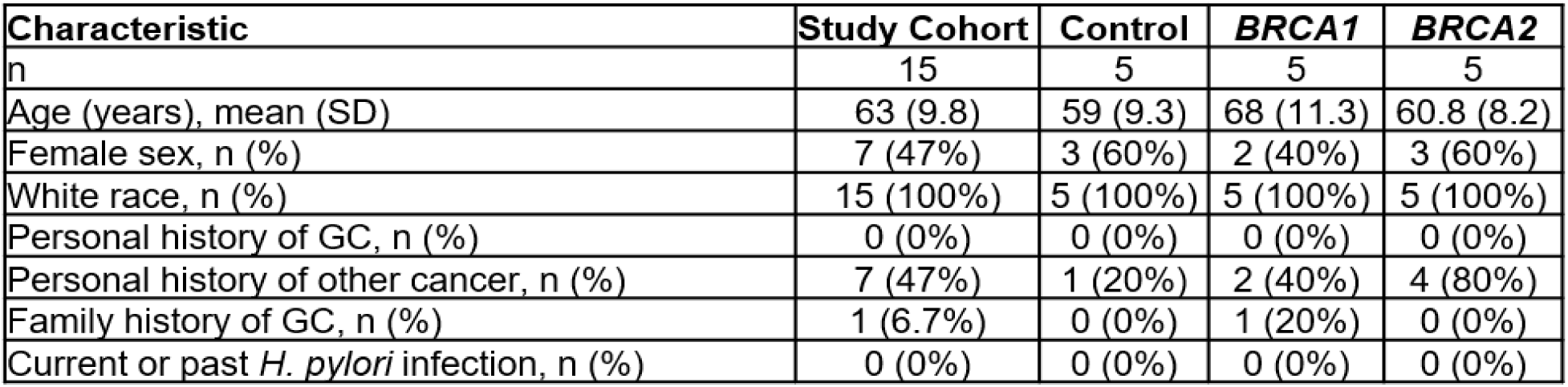
Baseline characteristics of the study cohort.

| Characteristic | Study Cohort | Control | *BRCA1* | *BRCA2* |
| --- | --- | --- | --- | --- |
| n | 15 | 5 | 5 | 5 |
| Age (years), mean (SD) | 63 (9.8) | 59 (9.3) | 68 (11.3) | 60.8 (8.2) |
| Female sex, n (%) | 7 (47%) | 3 (60%) | 2 (40%) | 3 (60%) |
| White race, n (%) | 15 (100%) | 5 (100%) | 5 (100%) | 5 (100%) |
| Personal history of GC, n (%) | 0 (0%) | 0 (0%) | 0 (0%) | 0 (0%) |
| Personal history of other cancer, n (%) | 7 (47%) | 1 (20%) | 2 (40%) | 4 (80%) |
| Family history of GC, n (%) | 1 (6.7%) | 0 (0%) | 1 (20%) | 0 (0%) |
| Current or past *H. pylori* infection, n (%) | 0 (0%) | 0 (0%) | 0 (0%) | 0 (0%) |

**Table S2:**
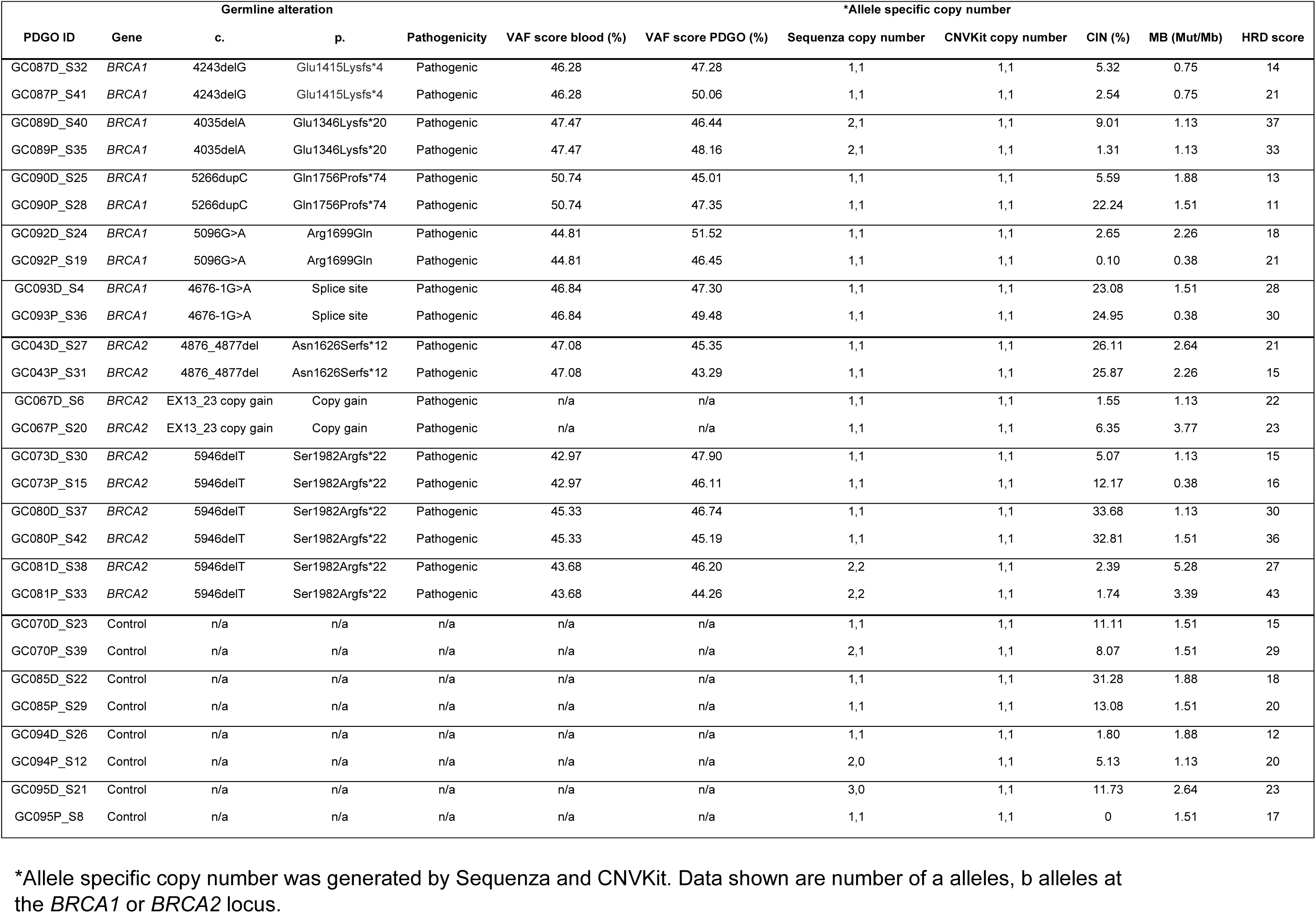
Individual PDGO DNA sequencing data.

